# Performances of targeted RNA-sequencing for the analysis of fusion transcripts, gene mutation and expression in haematological malignancies

**DOI:** 10.1101/2020.09.07.285981

**Authors:** Sandrine Hayette, Béatrice Grange, Maxime Vallee, Claire Bardel, Sarah Huet, Isabelle Mosnier, Kaddour Chabane, Thomas Simonet, Marie Balsat, Maël Heiblig, Isabelle Tigaud, Franck E. Nicolini, Sylvain Mareschal, Gilles Salles, Pierre Sujobert

## Abstract

RNA sequencing holds great promise to improve the diagnostic of haematological malignancies, because this technique enables to detect fusion transcripts, to look for somatic mutations in oncogenes, and to capture transcriptomic signatures nosological entities. However, the analytical performances of targeted RNA sequencing have not been extensively described in diagnostic samples. Using a targeted panel of 1385 cancer-related genes in a series of 100 diagnosis samples and 8 controls, we detected all the already known fusion transcripts, and also discovered unknown and/or unsuspected fusion transcripts in 12 samples. Regarding the analysis of transcriptomic profiles, we show that targeted RNA sequencing is performant to discriminate acute lymphoblastic leukemia entities driven by different oncogenic translocations. Additionally, we show that 86% of the mutations identified at the DNA level are also detectable at the mRNA level, except for nonsense mutations which are subjected to mRNA decay. We conclude that targeted RNA sequencing might improve the diagnosis of haematological malignancies. Standardization of the preanalytical steps and further refinements of the panel design and of the bioinformatical pipelines will be an important step towards its use in standard diagnostic procedures.

## Introduction

In haematological malignancies as in cancer in general, the goal of the diagnosis procedures is not only to accurately classify the patient’s disease according to the consensual WHO guidelines^1^, but also to identify biomarkers of prognostic or predictive values. A part of this information can be captured by morphology and immunophenotyping, but it relies more and more on the analysis of the genomic alterations of the neoplastic cells^2^. Nowadays, conventional cytogenetics and targeted sequencing of relevant genes are still the standard procedures. However, technological outbreaks such as whole genome sequencing, ATAC-sequencing or RNA sequencing (RNA-seq) might refine the diagnosis by unravelling genomic alterations outside coding regions^3^, epigenetic signatures^4,5^ and gene expression profiles respectively^6^.

In this study, we have chosen to assess the diagnostic value of RNA-seq, because this technique allows to explore three levels of genetic information: gene sequence, gene fusions, and gene expression. Interestingly, each of these different levels of analysis brings independent information about the neoplastic cell, and accordingly their integration should refine the precision of the diagnosis. For example, acute myeloid leukaemia (AML) patients prognosis is evaluated by cytogenetics (copy number abnormalities and structural variants), further refined by the analysis of the mutational status of a few genes, and could maybe be improved by transcriptomic signatures such as the LSC17 which is a proxy of the number of leukemic stem cells^7^.

Different techniques of library preparation for RNA-seq have been described, enabling the analysis of all the RNA molecules of a sample, or using enrichment step to target genes of interest such as messenger RNA, or small RNA species. Of note, the choice of the library preparation should optimize the balance between the number of targets of interest and the required depth of sequencing, in order to remain economically affordable in a routine setting. To date, most of the genes involved in cancer have been already identified by large programs of whole exome sequencing^8^. Based on these considerations, we have decided to evaluate the performances of a targeted RNA-seq panel of 1385 genes involved in cancer biology. We present here the analytical performances of targeted RNA-seq to detect fusion transcripts, to identify transcriptional profiles associated with clinically relevant entities, and to detect the recurrent mutations with clinical significance in haematological malignancies.

## Material and methods

### Samples

One hundred diagnosis samples from patients with the following haematological malignancies were included: acute leukaemia (acute myeloid leukaemia (AML), n=51 including 7 acute promyelocytic leukaemias (APL), B-cell acute lymphoblastic leukaemia (B-ALL, n=27), mixed phenotype acute leukaemia (n=1), T-cell acute lymphoblastic leukaemia (T-ALL, n=1)), myeloproliferative neoplasms (chronic myeloid leukaemia (CML, n=12) and other myeloproliferative neoplasms (n=2)), hypereosinophilic syndromes (HES, n=3)), chronic myelomonocytic leukemia (CMML, n=2) and myelodysplastic syndrome with multilineage dysplasia (n=1). These samples were chosen to enrich the cohort in fusion transcript due to chromosomal translocations, based on the results of conventional cytogenetics, in order to test the performances of targeted RNA-seq to detect fusion transcripts. Moreover, we used four controls (C1 to C4) prepared by pooling blood samples from 5 healthy donors for each, and four bone marrow samples from healthy donors. The characteristics of the samples are provided in supplemental Table 1. The procedures followed were in accordance with the Helsinki declaration, as revised in 2008.

Cytogenetic R and G-banding analyses were performed according to standard methods. The definition of a cytogenetic clone and description of karyotypes followed the current International System for Human Cytogenetic Nomenclature.

For a subset of samples (n=45), the analysis of a panel of 105 genes was already performed for routine diagnostic procedures, as already described^9^.

### RNA extraction

Three different protocols of RNA extraction were used (supplemental Table 1). For ALL samples and 3 bone marrow samples from healthy donors, RNA was extracted with Trizol reagent (TRIZ: Invitrogen, Carlsbad, CA, USA). For AML samples RNA was extracted with NucleoSpin RNA kit (MN: Macherey Nagel, Düren, Germany). For CML, HES, CMML and MDS samples, RNA was extracted with MN or the Maxwell 16LEV simplyRNA Blood Kit (Max: Promega, Madison, WI, USA). For control samples C1 to C4, RNA was extracted after Ficoll enrichment with either Trizol or MN methods, in order to assess the effect of extraction protocol on transcriptomic analysis performances.

### RNA sequencing

Library preparation for NGS sequencing was performed using TruSight RNA Pan-Cancer Panel (Illumina, SanDiego, CA, USA) targeting 1385 genes involved in cancer biology. Libraries from 16 samples were multiplexed and sequenced on a Nextseq 500 device with a 2×81 paired-end run on a mid-output flowcell according to the manufacturer’s instructions (mean number of reads by sample: 32.10^6^; range 20 to 59.10^6^).

### Bioinformatical analysis

After demultiplexing, adapter sequences were trimmed with Cutadapt and reads were mapped to the human genome (GRCh37). The detection of gene fusions was performed first with the commonly used STAR-Fusion pipeline (parameters: --FusionInspector validate) and STAR-2pass^10^, and all the negative samples were reanalysed with the recently launched nf-Core^11^ and Arriba (https://github.com/suhrig/arriba/) pipelines. Putative fusions were validated by reverse transcription and polymerase chain reaction (primers sequences are provided in supplemental Table 2). Gene expression analysis (after trimmed mean of M values (TMM) normalization), principal Component Analysis (PCA), k-means clustering, two tailed t-test and Heat Map generation followed by hierarchical clustering were performed using Omics Explorer software (Qlucore AB, Lund, Sweden). For gene mutation analysis on RNAseq data, we looked at all the mutations found at the DNA level by combining the same homemade workflow as for DNA, and visual inspection of the BAM files in case of unfound mutation. In brief we first gather the variant alleles called with Freebayes and Varscan2^12,13^. Among this raw set, we kept alleles whose read frequency was either above 20%, or for those below, if their frequency was more than 5 fold the median of the frequencies of all the samples from the same run. A second filtering step was applied to get rid of variants whose occurrence was above 1% in GnomAD mixed populations^14^.

## Results

### Identification of fusion transcripts

Fusion transcript positivity threshold was determined by the detection of at least one junction read and one spanning read between two different genes. All putative new fusion transcripts have been validated by PCR. All of the 57 rearrangements identified by cytogenetics or molecular biology were identified by targeted RNA-seq (Figure 1). Notably, RNA-seq detected all the *BCR-ABL1* canonical and rare transcripts (e13a2 (n=2); e14a2 (n=5); e1a2 (n=4); e1a3 (n=2); e6a2 (n=1); e13a3 (n=3); and e19a2(n=2)), as well as all the *PML-RARA* transcripts (BCR1 (n=2) BCR2 (n=2); BCR3(n=3)) and *MLL* (*KMT2A*) fusions (n=19). Of note, two samples with *FIP1L1-PDGFRA* fusion transcripts and one with *KMT2A* duplication were missed when analysed with the STAR-fusion pipeline, but recovered with nf-core and Arriba bioinformatics pipelines.

**Figure 1:**
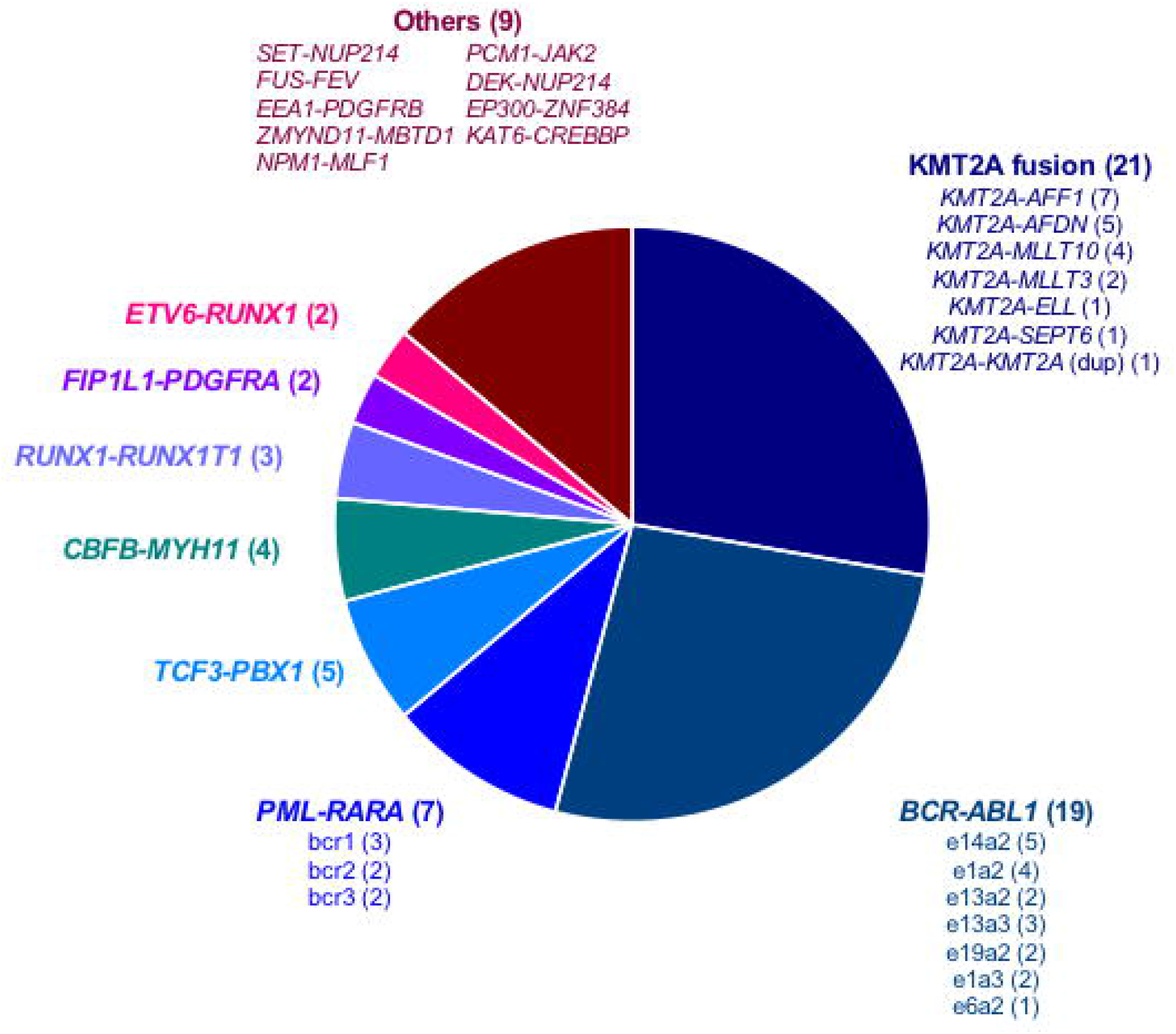
Description of the 72 fusion transcripts detected by targeted-RNAseq in the whole cohort.

Eighteen samples had a chromosomal translocation without detected fusion transcript based on routine molecular biology. In 5 patients, targeted RNA-seq identified a fusion transcript already described in the literature (*KAT6-CREBP, NPM1-MLF1, PCM1-JAK2, DEK-NUP214, ZMYND11-MBTD1*^15^) (Figure 1). In two additional patients, a new fusion transcript was identified and confirmed by RT-PCR and Sanger sequencing (*FUS-FEV; EEA1-PDGFRB*). These fusion transcript were in frame, probably leading to the expression of an abnormal fusion protein (Figure 2A and B). No evidence of fusion transcript was found in the 11 remaining samples.

**Figure 2:**
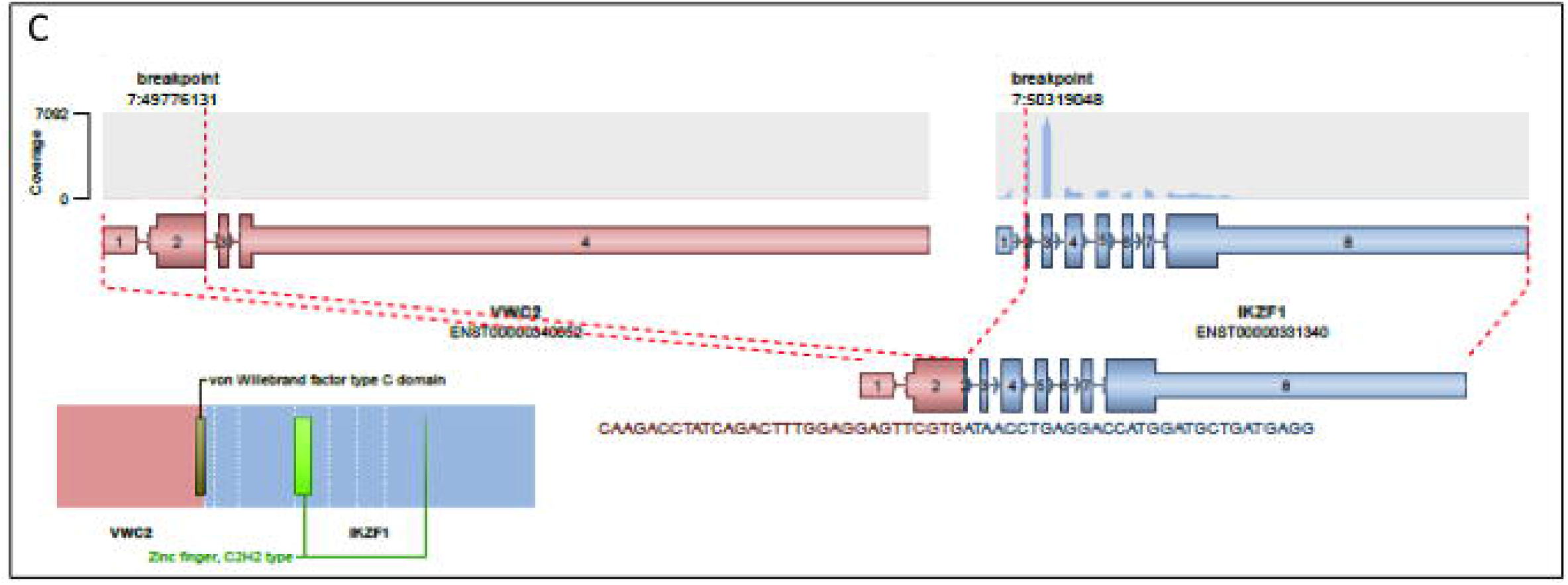
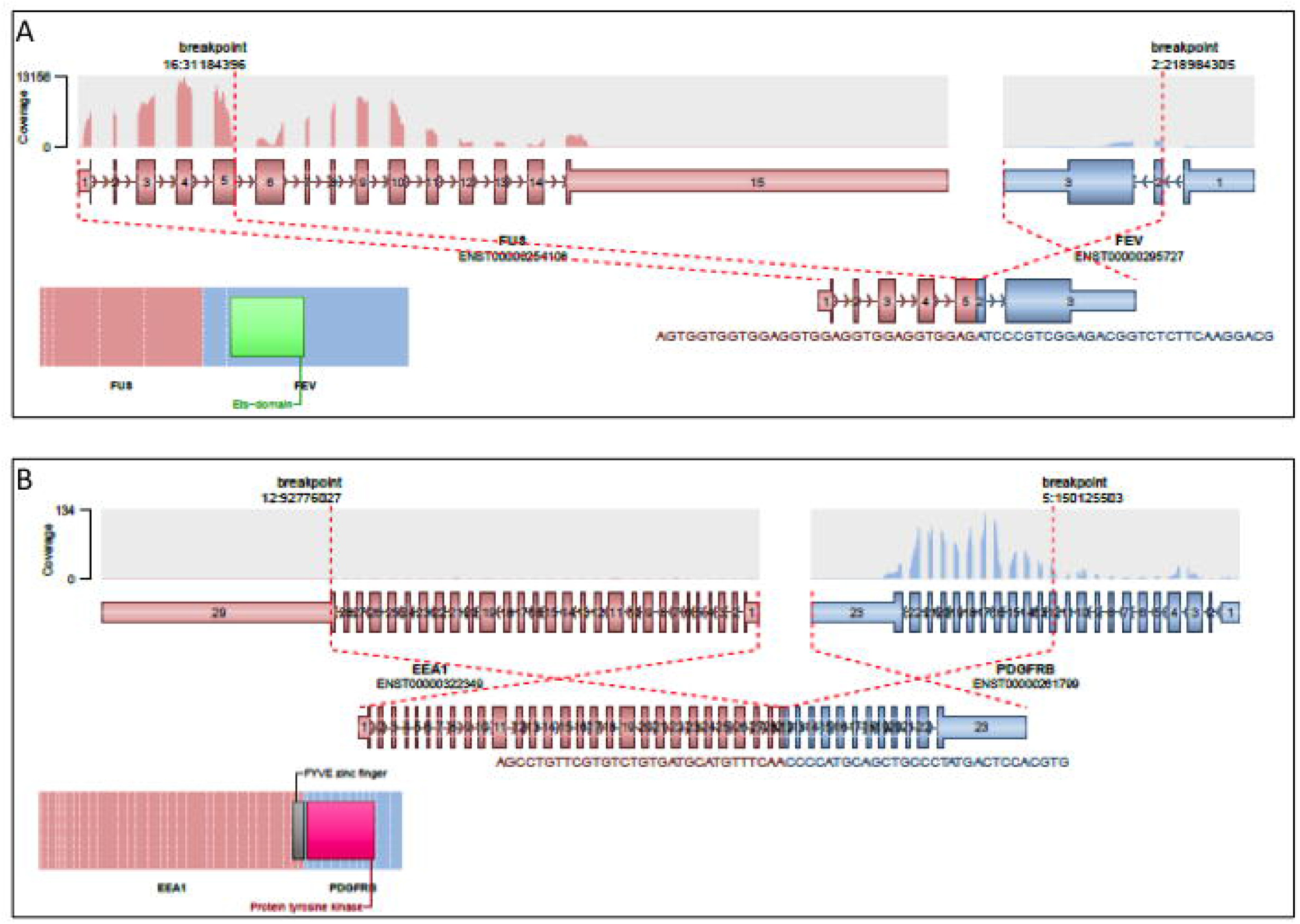
Description of the three new fusion transcripts discovered in this cohort. Schematic representation of the 3 new fusion transcripts identified by targeted RNAseq : *FUS-FEV* from t(2;16) (A), *EEA1-PDGFRB* from t(5;12) (B) and *VWC2-IKZF1* (C). For each fusion transcript is provided a schematic representation of the translocation at the genomic level, a graphical representation of the coverage depth in the targeted RNA-seq, and a schematic representation of the protein fusion.

Finally, we detected a fusion transcript in five samples without detectable translocation on conventional cytogenetics: *SET-NUP214, EP300-ZNF384, KMT2A-MLLT4, KMT2A-MLLT10, VWC2-IKZF1* (Figure 1). The *VWC2-IKZF1* fusion transcript (Figure 2C), never described so far, was detected in an acute lymphoblastic leukemia with a t(9;22) leading to the expression of the *BCR-ABL1*-transcript (patient 10, supplemental data Table 1). We hypothesize that this fusion might represent a new mechanism of *IKZF1* gene inactivation recurrently identified in Phi+-ALL^16,17^.

As it was previously described in non-cancer tissues and cells^18^, several fusions with open reading frame were also detected in control and patients’ samples. Some of them such as *TFG-GPR128, POLE-FUS* or *OAZ1-DOT1* were expressed at high level and have been also validated by RT-PCR and sequencing.

Finally, in order to assess the sensitivity threshold of RNA-seq to detect fusion transcripts, we analysed serial dilutions of two patients with *PML-RARA* and *BCR-ABL* fusion transcripts respectively. The detection threshold was below 6% for both fusion transcripts.

### Transcriptome analysis

The analysis of transcriptome in the routine diagnosis procedure is technically challenging, because of interferences linked to the source of the samples analysed (e.g. bone marrow vs peripheral blood), the preparation of the samples (isolation of the mononucleated cells with Ficoll or not), the RNA extraction method, and the batch effect of library preparation and sequencing. Instead were developed signatures based on a limited number of transcripts analysed with technical platforms such as RT-MLPA^19^ or Nanostring technology^7,20^. Here we assessed the feasibility of transcriptome analysis based on RNA-seq of a panel of 1385 genes.

First, we evaluated the magnitude of systematic biases in transcriptomic analysis introduced by the protocol of RNA extraction and the sequencing process. The same blood samples from healthy donors were extracted after Ficoll enrichment either with Trizol (n=4), or with Macherey Nagel kits (n=4). A supervised analysis based on extraction method identified 20 differentially expressed genes (fold change threshold 2, false discovery rate q<0,05) (Figure 3A). On the contrary, when we compared the transcriptome of RNA extracted from blood samples from healthy donors, whose RNA-seq libraries and sequencing were not prepared and run the same day, there was no gene differentially expressed according to the batch of library preparation or sequencing (fold change threshold 2, false discovery rate q <0,05, data not shown).

**Figure 3:**
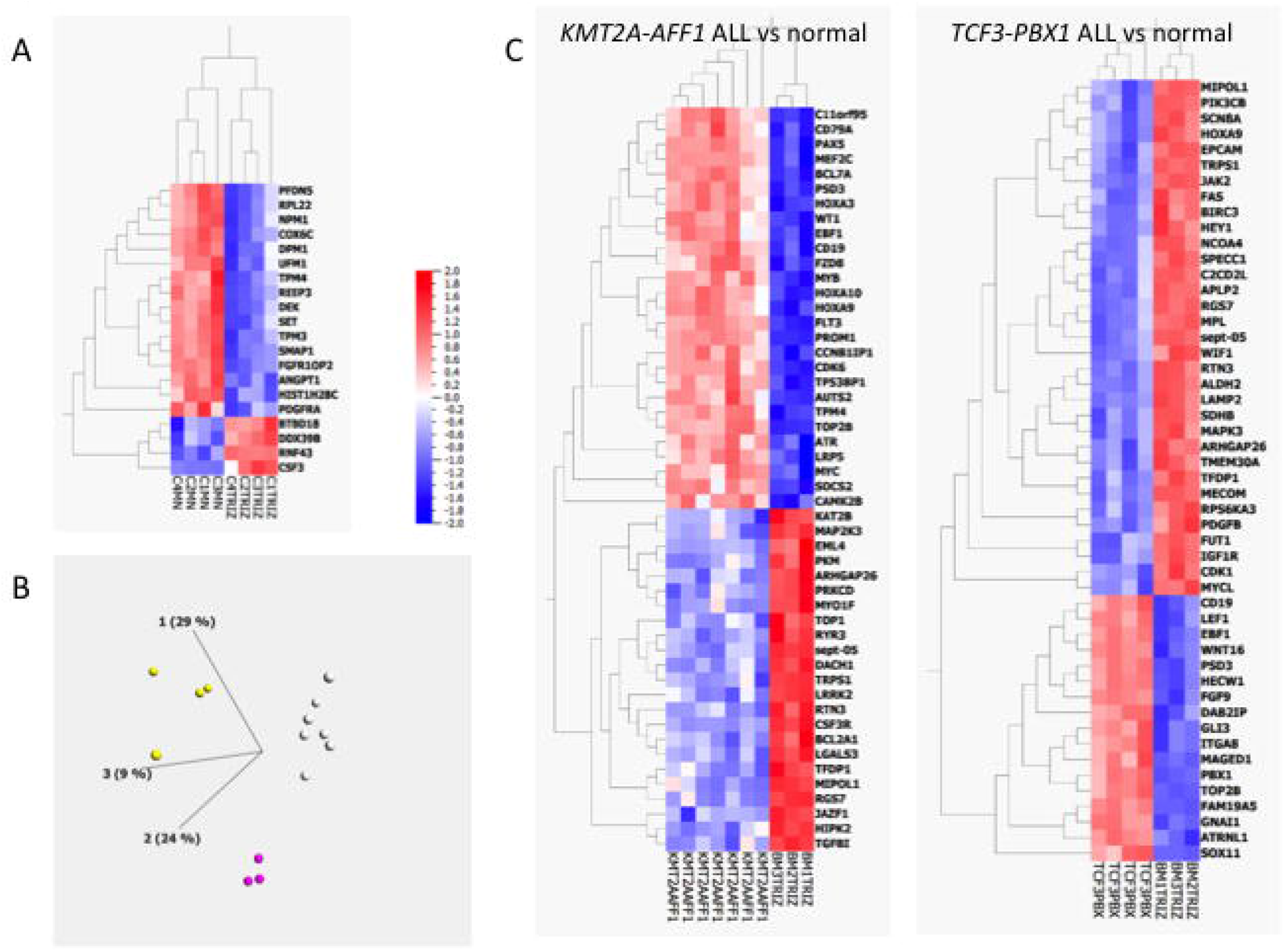
A: Heatmap representation of the 20 genes differentially expressed (fold change >2, q value <0,05) between the same control blood samples after RNA extraction with two different methods (trizol vs Macherey Nagel). B: Principal component analysis and unsupervised k-means clustering of 14 samples processed with the same preanalytical steps (*KMT2A-AFF1* ALL, white dots, n=7, *TCF3-PBX1* ALL, yellow dots, n=4, normal bone marrow samples, purple dots, n=3). C: Heatmaps showing the 50 most differentially expressed genes between normal samples and *KMT2A-AFF1* ALL samples (left) and between normal samples and *TCF3-PBX1* ALL samples (right).

Then, we analysed bone marrow samples extracted with the same method (Trizol) from three groups with at least 3 patients: ALL with *KMT2A-AFF1* (n=7), ALL with *TCF3-PBX1* (n=4), and normal bone marrow controls (n=3). Of note, these RNA were extracted at the time of diagnosis, over a period of 19 years, introducing a potential bias due to differences in RNA conservation. Clustering of these samples in three categories (by the k-means method) distinguishes the three groups of samples according to the diagnosis, with no misclassification (Figure 3B). The analysis of the 50 most differentially expressed genes between control and both type of ALL confirmed previously described features such as *HOXA3, HOXA9, HOXA10* and *FLT3* overexpression in *KMT2A-AFF1*^21^ and *CD19, WNT16* and *PBX1* up-regulation in *TCF3-PBX1* (Figure 3C)^22^.

### Detection of gene mutations

Forty five patients analysed with targeted RNA-seq were also analysed at the DNA level for a panel of 105 genes recurrently mutated in haematological malignancies^9^. Among the 95 genes captured in both panels, 122 mutations were detected at the DNA level in 39 different genes. As shown in Figure 4, 106/122 (87%) mutations identified at the DNA level were also found in the RNA seq data. Among the 16 mutations missed at the mRNA level, frameshift mutations were overrepresented (missed mutations 11/16 vs 12/106, Fischer’s exact test p<0.0001). Two other missed mutations (I1897T and G218V from *TET2* and *U2AF1*, respectively) were in low coverage areas (<30x).

**Figure 4:**
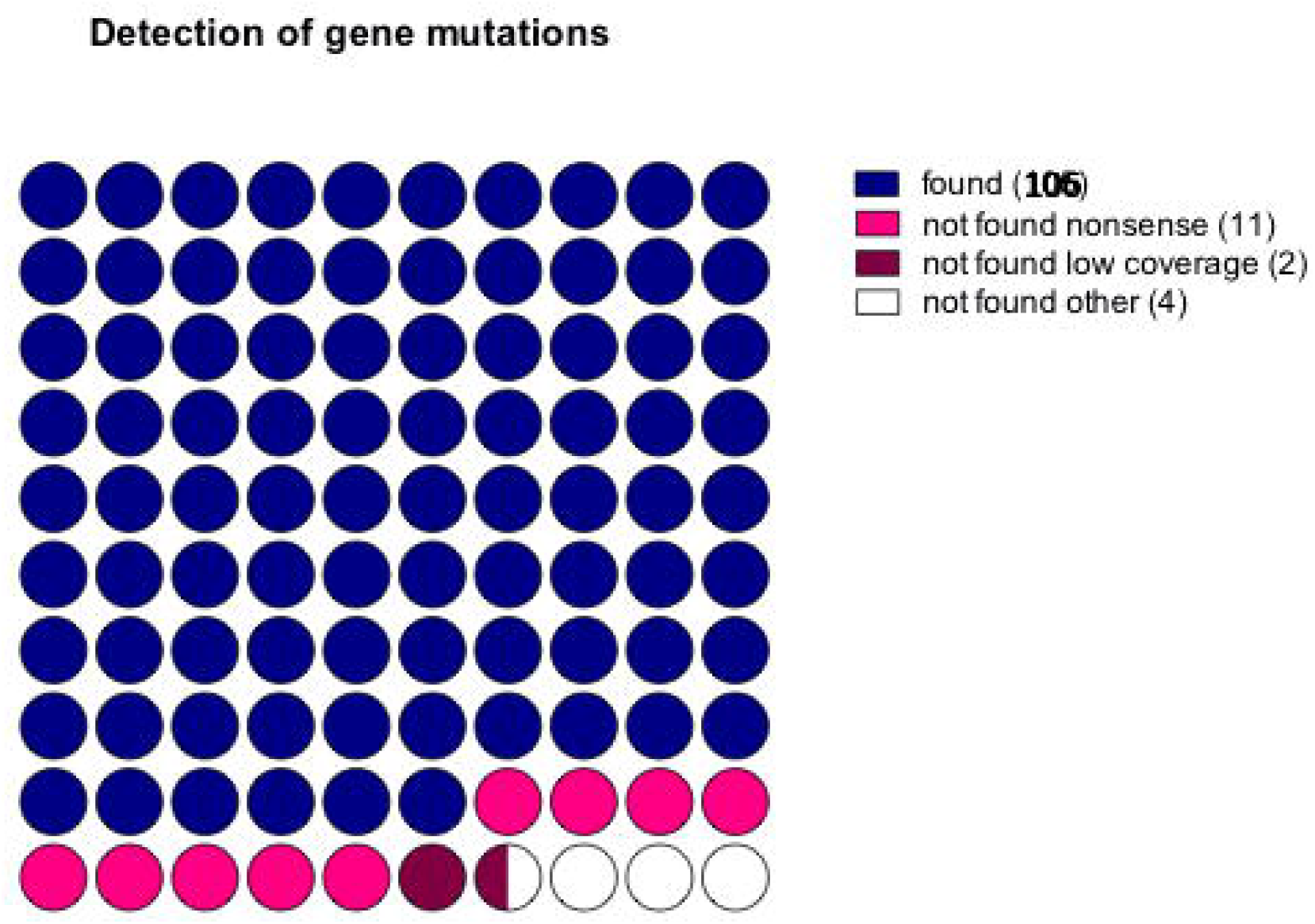
Performances of somatic mutations detection based on RNA-seq analysis. The relative number of mutations correctly identified or undetected (non-sense, low coverage or other) are presented.

## Discussion

This work reports the analytical performances of RNA-sequencing of a panel of 1385 genes to improve the diagnosis of haematological malignancies, based on a series of 100 diagnosis samples and 8 controls.

Overall, this technique detect 100% of fusion transcripts of these samples, including *FIP1L1-PDGFRA* fusions which often require nested PCR to be identified because of low levels of expression^23^. Of note, two fusion transcripts were found only by alternative bioinformatics pipelines, which highlights the major impact of the bioinformatics analysis on the performances of targeted RNA-seq. This might explain suboptimal detection of *KMT2A* and *PDGFRA* fusions in previous studies^24^. Interestingly, RNA-seq allowed the identification of 12 fusion transcripts which were not suspected with usual analysis recommended in the diagnosis of haematological malignancies^25^. As more and more case reports describe successful opportunistic use of targeted therapies in patients with fusion transcripts^26–28^, the identification of unexpected fusion transcripts might offer interesting targets in relapsed/refractory patients. In the series reported here, the patient with an hypereosinophilic syndrome associated with the novel *EEA1-PDGFRB* fusion was treated with imatinib and achieved a durable haematological remission. Moreover, as translocations are most of the time drivers events which are stable during disease evolution^30^, they can be used to track minimal residual disease with high sensitivity RT-qPCR, and adapt therapeutic intensity accordingly. However, it remains to be determined if the prognostic impact of minimal residual disease described for CBF AML^31^, CML, or APL is also true for the less recurrent fusion transcripts.

Regarding the analysis of the transcriptomic profile, we show that targeted transcriptome analysis can be used for nosological purposes if the pre-analytical workflow is the same for the samples analysed. Larger series are needed to precise the performances of targeted RNA-seq to resolve this task. Another interesting question would be to assess the performances of targeted RNA-seq to measure clinically relevant signatures such as the LSC17^7^ or the more recently described pLSC6^32^ signatures in AML, but it will need an optimization of the design of the panel to capture all relevant mRNAs.

The third clinical interest of targeted RNA-seq assessed here is the detection of acquired somatic mutations. Even if most of the mutations identified at the DNA level were found in RNA-seq data, the nonsense mutations were rarely detected. This is probably at least in part due to the phenomenon of mRNA decay, which degrades preferentially truncated mRNA^33^, and this will remain a biological limitation of RNA-seq for mutation assessment. Finally, given the growing importance of clonal architecture analysis based on variant allele frequency (VAF) deconvolution^34^, we should keep in mind that the VAF measured at the mRNA level might not be good surrogate markers of clonal architecture, because it takes into account allelic expression bias.

Altogether, RNA-seq of a targeted panel of genes might improve the diagnosis of haematological malignancies, and highlight potential therapeutic targets. Some of the limitations of this technique might be resolved with the optimization of the panel design and the bioinformatics pipelines for haematological malignancies. However, because some limitations have a biological explanation, such as poor performances to detect non-sense mutations, RNA-seq should not replace the analysis of genomic DNA, but could be rather a good orthogonal method for verifying genomic mutations and a powerful complement to increase the molecular characterisation of haematologic malignancies at diagnosis.

---

**Supplemental Table 1:** Characteristics of the samples included in the study.

**Supplemental Table 2:** Sequences of the primers used to validate the fusion transcripts.

**Supplemental Table 3:** List of gene mutations identified by DNA sequencing which were analysed on RNA-seq data.

