## Supplemental data Table 2 for "Performances of targeted RNA-sequencing for the analysis of fusion transcripts, gene mutation and expression in haematological malignancies"

| **FUSIONS** | **Forward (5'-3')** | **Reverse (5'-3')** |
|---|---|---|
| *EEA1-PDGFRB* | TGCCGACAGTGTGGAAATATCTTCTGTG | GCACAAGCTGGTCCCGCGGCAGCTC |
| *FUS-FEV* | GTGCGCGGACATGGCCTCA | GTTCTTCCCGTCCTTGAAGAG |
| *VWC2-IKZF1* | GACGAGAGCGGCTTCGTGTA | GACATGTCTTGACCCTCATCAG |
| **NON PATHOGENIC FUSIONS** | **Forward (5'-3')** | **Reverse (5'-3')** |
| *OAZ1-DOT1* | GCCGCACCATGCCGCTCCTAAGCCT | TCTCCATAGCGAGCTTGAGATCCGG |
| *TFG-GPR128* | GATAGTTCTGACCTTTCCTTTG | GCCATTTTCCCAGGTTCCACCAT |
| *POLE-FUS* | GCGCGGATGGCGAGGCCAGCA | CTTAATAATACCAATCTGCTTGAAGTA |
