## Supplemental data Table 1 for "Performances of targeted RNA-sequencing for the analysis of fusion transcripts, gene mutation and expression in haematological malignancies"

| N° | sample | Extraction type | Diagnosis | Karyotype | Fusion | junction+spaning read |
|---|---|---|---|---|---|---|
| 1 | BM | Trizol | B-ALL | NA | *TCF3-PBX1* | 924 |
| 2 | PBL | Trizol | B-ALL | 45,XX,der(7)t(7;8)(p12-13;q21), -8 [11] / 46,XX [13]. nuc ish (MLLx2)[200] | *unknown* | / |
| 3 | BM | Trizol | B-ALL | 46,XX, t(4;11)(q21;q23) [1] | *KMT2A-AFF1* | 33 |
| 4 | BM | Trizol | B-ALL | 46,XX,t(4;11)(q21;q23)[5]/46,XX[20] | *KMT2A-AFF1* | 11 |
| 5 | BM | Trizol | B-ALL | 46,XY,t(4;11)(q21;q23)[18]/46,XY[2] | *KMT2A-AFF1* | 54 |
| 6 | BM | Trizol | B-ALL | 46,XX,der(19)t(1;19)(q23;p13)[15]/46,XX[5] | *TCF3-PBX1* | 138 |
| 7 | BM | Trizol | B-ALL | 46,XX,t(4;11)(q21;q23)[6]/47,sl1,+der(4)t(4;11)[7]/47,sl2, ?ins(9;22)(q34;?q12)[2]/ 46,XX[5] | *KMT2A-AFF1* | 24 |
| 8 | BM | Trizol | B-ALL | 46,XY,t(4;11)(q21;q23)[8]/46,XY[12] | *KMT2A-AFF1* | 241 |
| 9 | BM | Trizol | B-ALL | 47,XX,t(1;19)(q23;p13),add(12)(p11),+mar[5]/46,XX[15] nuc ish(MLLx2)[200/200],(BCR,ABL)x2[200/200] | *TCF3-PBX1* | 525 |
| 10 | BM | Trizol | B-ALL | 46,XY,t(9;22)(q34;q11)[20] nuc ish(MLLX2)(ABLX3,BCRX3,ABLconBCRX2)[196/200] | *BCR-ABL1 e1a3+/VWC2-IKZF1* | 176_119 |
| 11 | BM | Trizol | B-ALL | 45,XX,t(9;22)(q34;q11),-10,del(11)(q12q24)[14]/45,XX,t(9;22)(q34;q11),add(9)(q24), -11[4]/46,XX[2] nuc ish(MLLX1)[154/200],(ABLX3,BCRX3,ABLconBCRX2)[96/200],amp(BCRconABL)[88/200] | *BCR-ABL1 e1a2* | 918 |
| 12 | PBL | MN | B-ALL | 46,XY,t(9;22)(q34;q11)[6]/45,XY,der(9)(13qter->13q12::9p12->9q34::22q11->22qter),-13,der(22)t(9;22)(q34;q11)[12] | *BCR-ABL1 e1a2* | 239 |
| 13 | BM | Trizol | B-ALL | 46,XX,dup(1)(q11q34),der(19)t(1;19)(q23;p13)[12]/46,XX[8] nuc ish(MLLX2),(ABL,BCR)X2[200/200] | *TCF3-PBX1* | 599 |
| 14 | BM | Trizol | B-ALL | 46,XX[20] nuc ish(ABL,BCR)X2[200/200],(MLLX2)[200/200],(TELX2,AMLX2,TELconAML1X1)[152/200] | *ETV6-RUNX1* | 230 |
| 15 | BM | Trizol | B-ALL | 46,XY,t(9;22)(q34;q11)[1]/47,idem,del(6)(q21q26),+10,add(14)(q32),dic(1;20)(p11;q11)[19] | *BCR-ABL1 e14a2* | 274 |
| 16 | PBL | Trizol | B-ALL | 46,XX,add(6)(q26),add(8)(q21),del(9)(q21q33),der(19)t(1;19)(q22;p13.3).ish(3' TCF3+,5'TCF3+)[5]/46,XX,add(6)(q27),del(11)(q22)[2]/46,XX[13] | *TCF3-PBX1* | 70 |
| 17 | BM | Trizol | B-ALL | 47,XY,del(6)(q21),+21[7]/49,idem,+16,+19[6]/85-87,<4n>,XXY,-1,-3,del(6)(q21)X2,-7,-13,+mar,inc[cp6]/46,XY[1] nuc ish(ETVX4,RUNX1X5,ETV6conRUNX1X2)[80/100],(ABLX4,BCRX4)[81/100],(KMT2AX3)[38/100] | *ETV6-RUNX1* | 357 |
| 18 | BM | Trizol | B-ALL | 47,XX,t(4;11)(q21;q23),+6[19]/46,XX[1] nuc ish(KMT2AX2,5'KMT2Asep3'KMT2AX1)[90/100],(ABL,BCR)X2[200/200] | *KMT2A-AFF1* | 47 |
| 19 | BM | Trizol | B-ALL | 44,XY,-4,add(14)?t(7;14)(q21;q32),del(5)(p15),i(17)(q10),-15,-18,+mar[7]/75,XXY,idemX2[1]/46,XY[13].nuc ish(KMT2AX2)[200/200],(BCRX3,ABLX2)[10/200],(CMYCX2)[200/200] | *NO* | / |
| 20 | BM | Trizol | B-ALL | 46,XY,del(20)(q11q13)[14]/46,idem,t(9;11)(q26;q23)[6] nuc ish(KMT2AX2)[192/200] | *NO* | / |
| 21 | PBL | Trizol | T-ALL | 49,XY,+Y,del(1)(q31q43),+8,+21[3]/46,XY[17] | *unknown* | / |
| 22 | BM | Trizol | B-ALL | 46,XY,add(9)(p23)[8] /46,XY[12] nuc ish(ABL,BCR)X4[35/100],(KMT2AX4)[28/100],(FGRF1X4)(20/200],(PDGFRBX4)[20/200] | *NO* | / |
| 23 | BM | Trizol | B-ALL | 46,XX[20] nuc ish(KMT2AX2),(ABL,BCR)X2[200/200] | *EP300-ZNF384* | 242 |
| 24 | PBL | Trizol | B-ALL | 48,XX,+X,+21[10]/46,XX(10] nuc ish(KMT2AX2)[200/200],(BCR,ABL)X2[200/200] | *NO* | / |
| 25 | BM | Trizol | B-ALL | 46,XX,t(4;11)(q21;q23)[18]/46,XX[2] .nuc ish(KMT2AX2,5'KMT2Asep3'KMT2AX1)[92/100],(ABL,BCR)X2,(ETV6,RUNX1)X2[200/200] | *KMT2A-AFF1* | 240 |
| 26 | BM | trizol | B-ALL | 47,XX,del(4)(q13q35),del(8)(q12q24),add(14)(q23)?der(14)t(1;14)(q24;q23),+21[3]/46,XX[10] .nuc ish(IgH, BCR, ABL1, KMT2A, ETV6)X2,(RUNX1X3)[30/100] | *NO* | / |
| 27 | BM | trizol | B-ALL | 58,XY,+X,+Y,dup(1)(q21q34),+4,+5,+add(6)(q27),+10,+14,+17,+18(X2),+21(X2)[6]/46,XY[4].nuc ish(ABL1,KMT2A,BCR)X2[200] | *NO* | / |
| 28 | mo | trizol | B-ALL | 46,XY,der(3)(?::3p21->3q12::?::3q12->3q26::?::3q26->3qter),add(14)(q32)[20].nuc ish(IgHcX2,IgHVdimX1)(IgHCconIgHVdimX1)[96/100],(MECOM,BCL6)X2)[200] | *NO* | / |
| 29 | BM | MN | APL | 46,XY, t(15;17)(q22;q21) [15]/46,XY[5] | *PML-RARA bcr2* | 51 |
| 30 | PBL | MN | APL | 46,XX,t(15;17)(q22;q12)[20]nuc ish(RARAX2,5'RARAsep3'RARAX1)[196/200](MLLX2)[200/200] | *PML-RARA bcr2* | 287 |
| 31 | PBL | MN | APL | 46,XY,t(15;17)(q12;q21)[15] | *PML-RARA bcr1* | 19 |
| 32 | BM | MN | APL | 46,XX,t(4;22)(p15;q12),del(15)(q25),t(15;17)(q22;q21)[20] nuc ish(PMLX3,RARAX3,PMLconRARAX2)[83/100] | *PML-RARA bcr1* | 94 |
| 33 | PBL | MN | APL | 46,XX,t(15;17)(q24;q21)[17]/46,XX[3] nuc ish(PMLX3,RARAX3,PMLconRARAX2)[166/200],(MLLX2)[200/200] | *PML-RARA bcr3* | 73 |
| 34 | BM | MN | APL | 46,XX,t(15;17)(q22;q21)[18]/46,XX[2] nuc ish(PML,RARA)X2(PMLconRARA)X1[160/200] | *PML-RARA bcr3* | 73 |

| | | | | | | |
|---|---|---|---|---|---|---|
| 35 | BM | MN | APL | 46,XX,t(15;17)(q22;q21)[18]/46,XX[2] nuc ish(PMLX3,RARAX3,PMLconRARAX2)[196/200],(KMT2AX2)[200/200] | *PML-RARA bcr3* | 77 |
| 36 | PBL | Trizol | AML | 46,XY,t(12;19)(q21;q11)[34] | *unknown* | / |
| 37 | BM | Trizol | AML | 46,XY[20] | *NO* | / |
| 38 | BM | MN | AML | 46,XX,t(8;16)(p11;p13)[15]/46,XX[5] | *KAT6-CREBBP* | 35 |
| 39 | BM | MN | AML | 46,XX,t(3;5)(q23;q34)[15]/ 46,XX[5] | *NPM1-MLF1* | 15 |
| 40 | PBL | Trizol | AML | 46,XX,t(8;21)(q22;q22)[3]/46,XX[4] | *RUNX1-RUNX1T1* | 341 |
| 41 | BM | MN | AML | NA | *NO* | / |
| 42 | BM | Trizol | AML | 46,XY [20]. | *NO* | / |
| 43 | BM | MN | AML | 46,XX[20] | *KMT2A -MLLT10* | 21 |
| 44 | sg | | AML | 46-50,XY,del(5)(q13q32),del(7)(?q22q33),+8,add(21)(p12),-22,+1à2mars, +r[cp12]/ 46,XY[8] | *NO* | / |
| 45 | PBL | MN | AML | 46,XY,-11,+mar,inc[cp5] nuc ish(MLLX2)(5'MLLSep3'MLLX1)[110/200] | *KMT2A -MLLT10* | 12 |
| 46 | BM | MN | AML | 38-42,XY,-2,add(3)(p12),-5,del(6)(q16q27),-7,add(8)(p22),-9,-11, -15,-16,-17,-18,-22,+1-5mars[cp28] nuc ish(MLLX3)[76/200],(CBFBX1)[162/200] | *unknown* | 0 |
| 47 | BM | MN | AML | 46,XY,inv(16)(p13q22)[7]/46,XY[13] nuc ish(CBFBX2,5'CBFBsep3'CBFBX1)[170/200](MLLX2)[200/200] | *CBFB-MYH11* | 40 |
| 48 | mo | | AML | 47,XX,+21[4]/47,idem,der(16)t(1;16)(q22;q22)[19] nuc ish(MLLX2)[200/200] | *NO* | / |
| 49 | BM | MN | AML | 46,XY[20] nuc ish(MLLX2)[200/200] | *NO* | / |
| 50 | PBL | MN | AML | 46,XX,t(6;9)(p23;q34)[7]/46,XX[13] nuc ish(NUP214X3,DEKX3,DEKconNUP214X2)[40/100](MLLX2)(CBFB,MYH11X2)[200/200] | *DEK-NUP214* | 34 |
| 51 | mo | | AML | 47,XX,+4[20] nuc ish(MLLX2)(CBFB,MYH11)X2[200/200] | *NO* | / |
| 52 | BM | MN | AML | 46,XY,del(11)(q21q23)[7]/46,XY[13] nuc ish(MLLX2)[200/200] | *KMT2A-KMT2A (DUP)* | / |
| 53 | PBL | MN | AML | 46,XX,inv(16)(p13q22)[15]/45,sl,-X[14]/46,XX[4] nuc ish(CBFBX3,MYH11X3,CBFBconMYH11X2)[143/200] | *CBFB-MYH11* | 44 |
| 54 | BM | MN | AML | 46,XX,t(2;16)(q34;p12).ish(CBF+;MYH11+)[20] nuc ish (KMT2AX2)[200/200] | *unknown: FUS-FEV* | 57 |
| 55 | PBL | MN | AML | 92,XXYY,inv(16)(p13q22)X2[13]/46,XY[7] nuc ish(CBFBX6,MYH11X6,CBFBconMYH11X4)(KMT2AX4)[68/200] | *CBFB-MYH11* | 116 |
| 56 | BM | MN | AML | 46,XY,t(6;11)(q27;q23)[17]/XY[3] | *KMT2A -AFDN* | 101 |
| 57 | PBL | MN | AML | 46,XX,t(2;12)(q23;p11)[6]/45,idem,-X,add(8)(q23)[9] nuc ish(MLLX2)(AML1X3,ETOX3,AML1conETOX1)[90/100] | *RUNX1-RUNX1T1* | 490 |
| 58 | PBL | MN | AML | 46,XY,del(5)(q31q35),t(10;17)(p15;q22),add(12)(p13)[8]/46,XY[6].nuc ish(TELx2)[200/200],(PDGFRBx2)[200/200] | *ZMYND11-MBTD1* | 6 |
| 59 | BM | MN | AML | 46,XY[20] nuc ish(MLLX2)(5'MLLX3,3'MLLX2,5'MLLcon3'MLLX2)[31/100] | *KMT2A -AFDN* | 32 |
| 60 | BM | MN | AML | 46,XX,t(9;22)(q34;q11),inv(16)(p13q22)[20] nuc ish(MLLX2),(CBFBX3,MYH11X3,CBFBconMYH11X2)[92/100] | *BCR-ABL1 e14a2+CBFB-MYH11* | 73_67 |
| 61 | PBL | MN | AML | 46,XX,t(8;21)(q22;q22)[16]/46,XX[4] nuc ish(MLLX2)(CBFB,MYH11)X2,(AML1X3,ETOX3,AML1conETOX2)[96/100] | *RUNX1-RUNX1T1* | 221 |
| 62 | BM | MN | AML | 46,XY[20] nuc ish(KMT2AX2)[200/200] | *NO* | / |
| 63 | sg | MN | AML | 46,XX[20] nuc ish (KMT2AX2)[200/200],(PML,RARA)X2[200/200] | *NO* | / |
| 64 | mo | MN | AML | 46,XY[20] nuc ish(KMT2AX2)[200/200] | *NO* | / |
| 65 | BM | MN | AML | 46,XX,t(9;11)(p22;q23)[6]/46,XX[14] nuc ish(KMT2AX2,5'KMT2Asep3'KMT2AX1)[45/100] | *KMT2A -MLLT3* | 30 |
| 66 | PBL | MN | MPAL | 46-51,XX,+X,+4,del(5)(q23q34),+8,del(9)(q33q34),+10,+21[20] nuc ish(ABLX1,BCRX2)[92/100],(KMT2AX2)[200/200],(EGR1X1),(D5S23,D5S721)X3[98/100] | *SET-NUP214* | 192 |
| 67 | BM | MN | AML | 52,XX,t(6;11)(q26;q23),+der(6)t(6;11),+8,+14,+18,+19,+22[11]/53,idem,+dm[8]/46,XX[1] .ish t(6;11)(3'KMT2A+;5'KMT2A+),der(6)t(6;11)(3'KMT2A+) | *KMT2A -AFDN* | 59 |
| 68 | BM | MN | AML | 46,XX,der(21)(t(1;21)(q23;p13)[3]/46,XX[17] nuc ish(KMT2AX2)[192/200] | *unknown* | / |
| 69 | BM | MN | AML | 46,XY,t(X;11)(q22;q23)[15]/46,XY[5] nuc ish(MLLX2)(3'MLLX1,5'MLLX2)[90/100],(CBFB,MYH11)X2[200/200] | *KMT2A-SEPT6* | 109 |
| 70 | BM | MN | CMML | 46,XY,der(1)t(1;1)(p36;q21)[13]/46,XY[7] | *unknown* | / |
| 71 | BM | MN | AML | 46,XX,t(9;11)(p21;q23)[18]/46,XX[2] nuc ish(KMT2AX2)(5'KMT2Asep3'KMT2AX1)[60/100] | *KMT2A -MLLT3* | 26 |
| 72 | BM | MN | AML | 46,XY,t(9;11)(p21;q23)[4]/46,XY[16] nuc ish(KMT2AX2)(5'KMT2Asep3'KMT2AX1)[34/100] | *KMT2A -MLLT3* | 10 |

| | | | | | | |
|---|---|---|---|---|---|---|
| 73 | BM | MN | AML | 46,XX,t(6;11)(q27;q23)[15]/52,idem,+3,+4,+der(6)t(6;11),+8,+19,+21[5] nuc ish(KMT2AX2,5'KMT2Asep3'KMT2AX1)[67/100](KMT2AX2,5'KMT2AX2sep3'KMT2AX1)[24/100] | *KMT2A -AFDN* | 53 |
| 74 | BM | MN | AML | 46,XX,t(11;19)(q23;p13.1)[15]/46,XX[5].nuc ish(KMT2AX2)(5'KMT2Asep3'KMT2AX1)[56/100] | *KMT2A -ELL* | 55 |
| 75 | BM | MN | AML | 46,XY,inv(3)(q21q26),del(7)(q21q26)[3]/46,XY[5].nuc ish(KMT2AX2)[200/200],(MECOMX2,5'MECOM sep3'MECOMX1)(10/100],(D7Z1X2,D7S486X1)[10/100] | *NO* | / |
| 76 | BM | MN | AML | 46,XY,t(8;9)(p22;p24)[10]/46,XY[10] | *PCM1-JAK2* | 18 |
| 77 | BM | MN | AML | 45,XX,-7,der(10)(11pter->11p13::?::10p12->10qter),der(11)t(10;11)(p12;p13)[18]/46,XX[2].nuc ish(D7Z1,D7S486)X1[83/100],(5'KMT2AX3,3'KMT2AX2, 5'KMT2Acon3'KMT2AX2)[78/100](RUNX1,RUNX1T1,CBFB,MYH11)X2[20/200] | *KMT2A -MLLT10* | 5 |
| 78 | BM | MN | CMML | 46,XY,t(5;12)(q32;q23)[14]/46,XY[6].nuc ish(PDGFRBX2,5'PDGFRBsep3'PDGFRBX1)[129/200] | *unknown: EEA1-PDGFRB* | 7 |
| 79 | BM | MN | AML | 46,XY,t(3;12)(q26;q13),del(7)(q22)[1].nuc ish(MECOMX2,5'MECOMsep3'MECOMX1)[45/100],(D7Z1,D7S486)X1[51/100],(KMT2AX2)[200/200] | *unknown* | / |
| 80 | BM | MN | MDS | 46,XY,i(17)(q10)[1].nuc ish(PDGFRB, FIP1)X2[200/200] | *unknown* | / |
| 81 | BM | MN | AML | 46,XX,der(10)(10pter->10p14::11q12->11q23::10p12->10qter),der(11)(11pter->11q12::10p14->10p12::11q23->11qter)[11]/46,XX[2].nuc ish(KMT2AX2,5'KMT2Asep3'KMT2AX1)[91/100],(MLLT1,MLLT3,MLLT4,CBFB,MYH11,RUNX1,RUNX1T1)X2[200] | *KMT2A -MLLT10* | 151 |
| 82 | mo | | AML | 44,XY,dic(5;17)(q12;p11),del(7)(q22q36),dic(11;12)(p12;p12)[9].nuc ish(MECOM, RUNX1T1, KMT2A, CBFB, MYH11, RUNX1)X2[196/200] | *NO* | / |
| 83 | BM | Trizol | AML | 46,XX,t(6;20)(q22;q12)[11]/56-58,XX,+2,+4,+8,+9,t(9;22)(q34;q11),+10,+11,+1,+18,+21,+22,+1-2mars[9] | *unknown+ BCR-ABL1 e1a2* | 94 |
| 84 | BM | Trizol | CML | 46,XX,t(9;22)(q34;q11)[1]/ 47,idem,+8[20]/ 47,XX,+8[1]/ 46,XX[3] | *BCR-ABL1 e14a2* | 39 |
| 85 | PBL | Trizol | CML | 46, XX,t(9;22)(q34;q11),t(11;16)(p12;q21)[14]/ 46, XX[6] | *BCR-ABL1 e6a2* | 34 |
| 86 | sg | | SMP | NA | *NO* | / |
| 87 | PBL | Trizol | CML | 46,XX,t(9;22)(q34;q11)[20] | *BCR-ABL1 e1a3* | 40 |
| 88 | PBL | MN | CML | 46,XY,t(9;22)(q34;q11)[29]/47,idem,+8[3] | *BCR-ABL1 e13a3* | 74 |
| 89 | PBL | MN | CML | 46,XY,t(9;22)(q34;q11)[20] | *BCR-ABL1 e14a2* | 90 |
| 90 | PBL | MN | CML | 46,XX,t(9;22)(q34;q11)[20] | *BCR-ABL1 e14a2* | 36 |
| 91 | PBL | MN | CML | 46,XX,t(9;22)(q34;q11)[20] | *BCR-ABL1 e13a2* | 21 |
| 92 | PBL | Trizol | CML | 46,XX,t(9;22)(q34;q11)[20] | *BCR-ABL1 e13a3* | 50 |
| 93 | PBL | MN | CML | 46,XX,t(9;22)(q34;q11)[17]/47,idem,+der(22)t(9;22)(q34;q11)[3] | *BCR-ABL1 e19a2* | 48 |
| 94 | PBL | Maxwell | CML | 46,XY,t(9;22)(q34;q11)[20] | *BCR-ABL1 e13a2* | 45 |
| 95 | PBL | MN | CML | 46,XX,t(9;22)(q34;q11)[9]/47,idem,+8[11] | *BCR-ABL1 e19a2* | 23 |
| 96 | PBL | MN | SHE | 46,XY[20] nuc ish(PDGFR BX2)[200/200],(PDGFRalphaX2,CHIC2X1)[82/100] | *FIP1-PDGFRA* | 3 |
| 97 | PBL | MN | CML | 46,XY,t(9;22)(q34;q11)[16]/46,XY[4] | *BCR-ABL1 e13a3* | 39 |
| 98 | PBL | Maxwell | SHE | 46,XY[20] nuc ish(FIP1L1X2,CHIC2X1)[68/100] | *FIP1-PDGFRA* | 30 |
| 99 | PBL | Maxwell | SHE | 46,XY[20].nuc ish(PDGFR A, PDGFRB, FGFR1)X2[200/200],(20q12X1,20q13X2)[41/200] | *unknown* | / |
| 100 | PBL | Maxwell | SMP | 46,XY,t(12;14)(q22;q24)[20].nuc ish(FGFR1,ABL1,ETV6,BCR)X2[200] | *unknown* | / |
| 101 | PBL | MN | C1 | | *biais batch* | / |
| 102 | PBL | Trizol | C1 | | *biais batch* | / |
| 105 | PBL | MN | C2 | | *biais batch* | / |
| 106 | PBL | Trizol | C2 | | *biais batch* | / |
| 108 | PBL | MN | C3 | | *biais batch* | / |
| 109 | PBL | Trizol | C3 | | *biais batch* | / |
| 111 | PBL | MN | C4 | | *biais batch* | / |
| 112 | PBL | Trizol | C4 | | *biais batch* | / |

| | | | | | | |
|---|---|---|---|---|---|---|
| 113 | BM | Trizol | BM1 | | *biais batch* | / |
| 114 | BM | Trizol | BM2 | | *biais batch* | / |
| 115 | BM | Trizol | BM3 | | *biais batch* | / |
