## Supplemental data Table 3 for "Performances of targeted RNA-sequencing for the analysis of fusion transcripts, gene mutation and expression in haematological malignancies"

| *GENE* | *NM* | *Mutation* | *nature of the mutation* | *Detected by RNAseq* | *occurrence* |
|---|---|---|---|---|---|
| *ABL1* | NM_005157 | p.K247R | MAF<0.01 | Yes | 1 |
| *ABL1* | NM_005157 | p.S43N | MAF<0.01 | Yes | 1 |
| *ASXL1* | NM_015338 | p.G646Wfs*12 | fms | No | 2 |
| *ASXL1* | NM_015338 | p.K726fs | fms | Yes | 1 |
| *ASXL2* | NM_018263 | p.A497T | MAF<0.01 | Yes | 1 |
| *ASXL2* | NM_018263 | p.R591Pfs*18 | fms | No | 1 |
| *BCOR* | NM_001123385 | p.P678S | SNV | No | 1 |
| *BCOR* | NM_001123385 | p.V668* | fms | Yes | 1 |
| *CBL* | NM_005188 | p.R585C | SNV | Yes | 1 |
| *CBL* | NM_005188 | p.L380P | SNV | Yes | 1 |
| *CBL* | NM_005188 | p.C404W | SNV | Yes | 1 |
| *CBL* | NM_005188 | pR718* | fms | No | 1 |
| *CEBPA* | NM_001287424 | del85_309 | delfms | No | 1 |
| *CEBPA* | NM_001287424 | p.P23Nfs*81 | fms | No | 1 |
| *c-Kit* | NM_000222 | del416_421 | del_e8 | Yes | 1 |
| *c-Kit* | NM_000222 | p.N822K | SNV | Yes | 1 |
| *c-kit* | NM_000222 | p.D816V | SNV | Yes | 1 |
| *c-kit* | NM_000222 | p.D816Y | SNV | Yes | 1 |
| *c-Kit* | NM_000222 | p.Y418_D419insLP | SNV | Yes | 1 |
| *CREBBP* | NM_004380 | p.Y1204C | SNV | Yes | 1 |
| *CREBBP* | NM_004380 | p.M775I | SNV | Yes | 1 |
| *CREBBP* | NM_004380 | p.N327L | SNV | Yes | 1 |
| *CSF3R* | NM_156039 | p.E149D | MAF<0.01 | Yes | 1 |
| *CSF3R* | NM_156039 | p.N776* | fms | Yes | 1 |
| *CSF3R* | NM_156039 | p.R440N | SNV | Yes | 1 |
| *DNMT3A* | NM_175629 | p.A376T | SNV | Yes | 1 |
| *DNMT3A* | NM_175629 | p.F252V | SNV | Yes | 1 |
| *DNMT3A* | NM_175629 | p.R882H | SNV | Yes | 1 |
| *DNMT3A* | NM_175629 | p.N501S | SNV | Yes | 1 |
| *DNMT3A* | NM_175629 | p.T275Pfs*41 | fms | No | 1 |
| *ETV6* | NM_001987 | p.I318Tfs*20 | fms | No | 1 |
| *ETV6* | NM_001987 | p.I176T | SNV | Yes | 1 |
| *EZH2* | NM_004456 | p.G79R | SNV | Yes | 1 |
| *FBXW7* | NM_033632 | p.R465C | SNV | Yes | 1 |
| *FGFR1* | NM_001174067 | p.L292M | SNV | Yes | 1 |
| *FLT3* | NM_004119 | p.I836_M837insG | ins | Yes | 1 |
| *FLT3* | NM_004119 | p.I836del | del | Yes | 3 |
| *FLT3* | NM_004119 | p.D839G | SNV | Yes | 1 |
| *FLT3* | NM_004119 | p.D835V | SNV | Yes | 1 |
| *FLT3* | NM_004119 | p.D835Y | SNV | Yes | 1 |
| *FLT3* | NM_004119 | p.Y597_E598ins8 | ITD24bp | Yes | 2 |
| *FLT3* | NM_004119 | p.Q604_F605ins10 | ITD30bp | Yes | 1 |
| *FLT3* | NM_004119 | p.Y842C | SNV | Yes | 1 |
| *FLT3* | NM_004119 | p.D835E | SNV | Yes | 1 |
| *FLT3* | NM_004119 | p.N841I | SNV | Yes | 1 |
| *FLT3* | NM_004119 | p.M664V | SNV | Yes | 1 |
| *FLT3* | NM_004119 | p.N676K | SNV | Yes | 1 |
| *GATA2* | NM_001145661 | p.R398T | SNV | Yes | 1 |

| | | | | | |
|---|---|---|---|---|---|
| *GATA2* | NM_001145661 | p.P470R | SNV | Yes | 1 |
| *IDH1* | NM_005896 | p.R132S | SNV | Yes | 1 |
| *IDH1* | NM_005896 | p.R119W | MAF<0.01 | Yes | 1 |
| *IDH2* | NM_002168 | p.R140Q | SNV | Yes | 1 |
| *IDH2* | NM_002168 | p.P198A | MAF<0.01 | Yes | 1 |
| *IDH2* | NM_002168 | p.T435M | MAF<0.01 | Yes | 1 |
| *IDH2* | NM_002168 | p.R172K | SNV | Yes | 1 |
| *IKZF1* | NM_006060 | p.520N*29 | fms | Yes | 1 |
| *IKZF1* | NM_006060 | p.N159Y | SNV | Yes | 1 |
| *JAK2* | NM_004972 | p.V617F | SNV | Yes | 4 |
| *MPL* | NM_005373 | p.H624D | SNV | Yes | 1 |
| *NF1* | NM_001042492 | p.L2395Ffs*24 | fms | Yes | 1 |
| *NPM1* | NM_002520 | p.W288Cfs*12 | fms | Yes | 1 |
| *KRAS* | NM_033360 | p.G12V | SNV | Yes | 2 |
| *KRAS* | NM_033360 | p.G12D | SNV | Yes | 1 |
| *KRAS* | NM_033360 | p.G12V | SNV | No | 1 |
| *KRAS* | NM_033360 | p.L23R | SNV | Yes | 1 |
| *KRAS* | NM_033360 | p.G13D | SNV | Yes | 1 |
| *NRAS* | NM_033360 | p.G12S | SNV | Yes | 1 |
| *NRAS* | NM_002524 | p.G12V | SNV | No | 1 |
| *NRAS* | NM_002524 | p.Q61L | SNV | Yes | 1 |
| *PAX5* | NM_016734 | p.V26Afs*49 | fms | No | 1 |
| *PAX5* | NM_016734 | p.C64F | SNV | Yes | 1 |
| *PAX5* | NM_016734 | p.P80R | SNV | Yes | 1 |
| *PHF6* | NM_001015877 | p.R274N | SNV | Yes | 1 |
| *PHF6* | NM_001015877 | p.R335Mfs*15 | fms | Yes | 1 |
| *PTEN* | NM_000314 | p.A252T | SNV | Yes | 1 |
| *PTPN11* | NM_002834 | p.G503R | SNV | Yes | 1 |
| *RAD21* | NM_006265 | p.D276G | SNV | Yes | 1 |
| *RUNX1* | NM_001754 | p.N264fs | fms | Yes | 1 |
| *RUNX1* | NM_001754 | p.F330fs | fms | Yes | 1 |
| *RUNX1* | NM_001754 | p.A251Gfs*5 | fms | Yes | 1 |
| *RUNX1* | NM_001754 | p.D160N | SNV | Yes | 1 |
| *RUNX1* | NM_001754 | p.A149fs | SNV | Yes | 1 |
| *SETBP1* | NM_015559 | p.H1100R | SNV | Yes | 1 |
| *SETBP1* | NM_015559 | p.D868N | SNV | Yes | 2 |
| *SF3B1* | NM_012433 | p.R625C | SNV | Yes | 1 |
| *SRSF2* | NM_001195427 | p.P95H | SNV | Yes | 3 |
| *STAG2* | NM_001042749 | p.D1136Lfs*10 | fms | Yes | 1 |
| *TCF3* | NM_003200 | p.G385D | SNV | Yes | 1 |
| *TCF3* | NM_003200 | p.S514L | MAF<0.01 | Yes | 1 |
| *TCF3* | NM_003200 | p.S350fs ins 8nt | fms | No | 1 |
| *TCF3* | NM_003200 | p.G46R | SNV | Yes | 1 |
| *TCF3* | NM_003200 | p.H579R | SNV | Yes | 1 |
| *TET2* | NM_001127208 | p.R1167K | SNV | Yes | 1 |
| *TET2* | NM_001127208 | p.R1261H | SNV | Yes | 1 |
| *TET2* | NM_001127208 | p.V1718L | SNV | Yes | 1 |
| *TET2* | NM_001127208 | p.I1897T | SNV | No | 1 |
| *TET2* | NM_001127208 | p.V1718L | SNV | Yes | 2 |
| *TET2* | NM_001127208 | p.T625I | SNV | Yes | 1 |

| | | | | | |
|---|---|---|---|---|---|
| *TET2* | NM_001127208 | p.P1335dup | dup | Yes | 1 |
| *TET2* | NM_001127208 | p.A707Lfs*44 | fms | Yes | 1 |
| *TP53* | NM_000546 | p.Y236D | SNV | Yes | 1 |
| *TP53* | NM_000546 | p.G245S | SNV | Yes | 1 |
| *TP53* | NM_000546 | p.R282P | SNV | Yes | 1 |
| *TP53* | NM_000546 | p.A78V | neutral | Yes | 1 |
| *TP53* | NM_000546 | p.K132R | SNV | Yes | 1 |
| *TP53* | NM_000546 | p.R248N | SNV | Yes | 1 |
| *TP53* | NM_000546 | p.V157G | SNV | Yes | 1 |
| *TP53* | NM_000546 | p.R123* | fms | No | 1 |
| *U2AF1* | NM_006758 | p.S34F | SNV | Yes | 1 |
| *U2AF1* | NM_006758 | p.G218V | SNV | No | 1 |
